# How big should this object be? Perceptual influences on viewing-size preferences

**DOI:** 10.1101/2021.08.12.456159

**Authors:** Yi-Chia Chen (陳鴨嘉), Arturo Deza, Talia Konkle

## Abstract

When viewing objects depicted in a frame, observers prefer to view large objects like cars in larger sizes and smaller objects like cups in smaller sizes. That is, the visual size of an object that “looks best” is linked to its typical physical size in the world. Why is this the case? One intuitive possibility is that these preferences are driven by semantic knowledge: For example, when we recognize a sofa, we access our knowledge about its real-world size, and this influences what size we prefer to view the sofa within a frame. However, might visual processing play a role in this phenomenon—that is, do visual features that are related to big and small objects look better at big and small visual sizes, respectively, even when observers do not have explicit access to semantic knowledge about the objects? To test this possibility, we used “texform” images, which are synthesized versions of recognizable objects, which critically retain local perceptual texture and coarse contour information, but are no longer explicitly recognizable. To test for visual size preferences, we first used a size adjustment task, and the results were equivocal. However, clear results were obtained using a two-interval forced choice task, in which each texform was presented at the preferred visual size of its corresponding original image, and a visual size slightly bigger or smaller. Observers consistently selected the texform presented at the canonical visual size as the more aesthetically pleasing one. An additional control experiment ruled out alternative explanations related to size priming effects. These results suggest that the preferred visual size of an object depends not only on explicit knowledge of its real-world size, but also can be evoked by mid-level visual features that systematically covary with an object’s real-world size.

**Highlights:**

- We prefer to view large objects like cars large, and small objects like cups small
- Intuitively, such preferences may be driven by our knowledge of object sizes
- We used unrecognizable texforms of objects that preserved mid-level visual features
- Similar viewing size preferences can be revealed with these texforms
- Such preferences thus arise not only from knowledge but also from visual processing

## 1. Introduction

One of the most frequent everyday activities we engage in is inspecting objects. When we detect a bird in a tree, find a box of snacks lying deep in the fridge, or spot a product in an aisle of a shopping mall, we gather more information about the object by getting closer to it and stopping at a proper distance to look at it. This idea that each object has an optimal viewing distance, and that perception draws us to move our bodies to the distance that balances between deficiency on one hand (too far away) and excess on the other (too close), has been highlighted by philosophers of perception (Merleau-Ponty, 1962; Kelly, 2010). By moving closer to or farther from an object, the observer can adjust the visual size that the object subtends in their visual field. Indeed, research has found that given a picture of an object, there is a systematic, or “canonical”, visual size at which the object “looks best” in, and, curiously, this visual size is linked to the physical size of the object: When viewing items with a bigger physical size (e.g., a car), we prefer to view them at a bigger visual size; and when viewing items with a smaller physical size (e.g., a cup), we prefer a smaller visual size (Konkle & Oliva, 2011; Linsen, Leyssen, Sammartino, & Palmer, 2011; see also Eckstein, Koehler, Welbourne, & Akbas, 2017).

### 1.1 Knowledge of physical sizes

What is the nature of these physical size representations that drive the systematic canonical visual sizes? A likely candidate is the rich real-world size knowledge we eagerly pick up as we experience the world, evident in toddlers and even infants (e.g., Granrud, Haake, & Yonas, 1985; Long, Moher, Carey, & Konkle, 2019a, 2019b; Sensoy, Culham, & Schwarzer, 2020; Yonas, Pettersen, & Granrud, 1982). We can clearly learn the physical sizes of objects from our own past sensory experience (e.g., the size of our favorite toy from childhood) and build this knowledge further by incorporating semantic knowledge (e.g., even if you have never seen a “ranchu” or a picture of one, if you learn that it is a kind of goldfish, you might then infer that it is roughly the size of a typical pet goldfish; see Chen, Lu, & Holyoak, 2014). Further, size knowledge can be completely abstracted from direct sensory experience; for example, we can represent and reason about the physical size of an atom, the earth, or even of a unicorn.

Interestingly, this knowledge of objects’ real-world sizes seems to influence how we spatially allocate our visual attention (Collegio, Nah, Scotti, & Shomstein, 2019), demonstrating an example of interaction between size knowledge and other aspects of cognition. Thus, one possible account of canonical visual size is that it arises as a consequence of our abstract physical size knowledge.

### 1.2 Perception of physical sizes

Interestingly, along with the rich size knowledge we have, our visual systems seem to maintain perceptual representations that distinguish objects of different physical sizes as well. This point is revealed behaviorally with several different methods: Visually searching for a picture of a big object (e.g., a building) among an array of pictures of small objects (e.g., a flashlight, a cap, etc.) is faster than when searching for the same picture of the big object among other pictures of big objects (e.g., a bed, a boat, etc.; note that the *visual* size of all the items in the array is the same; Long, Konkle, Cohen, & Alveraz, 2016; Long et al., 2019a). This result indicates that there are systematic perceptual differences between big and small objects (as classes), that can be used to speed up visual search processes. For example, bigger objects tend to be boxier with higher spatial frequency, and small objects tend to be curvier and smoother (Konkle, 2011; Long et al., 2016).

An even stronger case for these systematic *perceptual* differences among objects of different physical sizes comes from a line of work using “texform images”—these are distorted images synthesized from images of recognizable objects, which critically retain some perceptual texture and coarse contour, while “knocking out” the object identity (See Figure 1; Long et al., 2016; Deza, Chen, Long, & Konkle, 2019). The facilitated visual search for big among small objects (and vice versa) persists with texform images (Long et al., 2016): That is, texforms of big objects were faster to find among texforms of small objects than among texforms of big objects. These results further support the claim that there are systematic perceptual differences in the shape and texture of objects of different real-world sizes.

**Figure 1.**
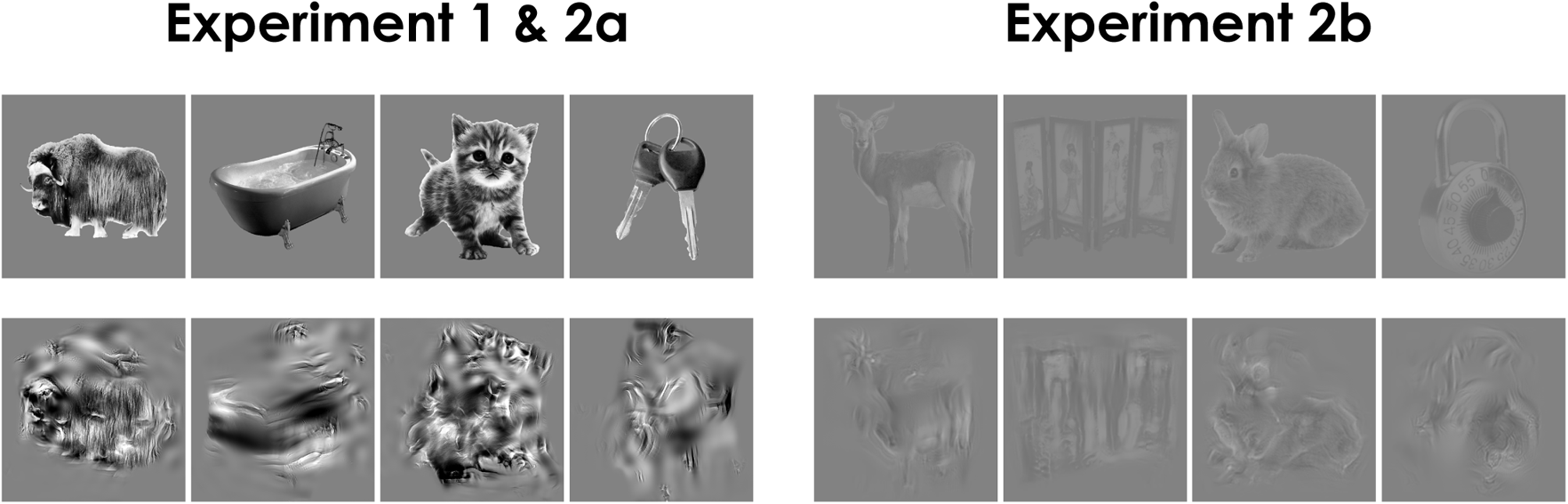
Example of intact and texform images from Experiment 1, 2a, and 2b.

These systematic perceptual differences between big and small objects not only influence visual search but also are powerful enough to interfere with even simple perceptual judgments about what image is bigger or smaller *on the screen*, a task that does not require any access to the identity or the real-world size of the objects. That is, people are faster to select the visually smaller of two objects on the screen if it is in fact smaller in the real world (Konkle & Oliva, 2011). Critically, the same effect was found using texforms (Long & Konkle, 2017; Long et al., 2019b): For example, people were faster to pick the visually smaller of two unrecognizable texforms, when the visually smaller texform was generated from a small object (e.g., key) than when the visually smaller texform was generated from a big object (e.g., piano). Thus, perceptual feature differences between big and small objects are sufficient to automatically influence visual size judgments. Finally, complementing these behavioral signatures, there is also evidence for different visual cortex sensitivity to these perceptual features: Different regions along the ventral stream respond more to big object texforms than small object texforms (and vice versa), with highly similar large-scale ventral stream topography as evoked when viewing recognizable objects with big vs small real-world sizes (Long, Yu, & Konkle, 2018).

Taken together, these studies prompt another possible account for canonical visual size: The preferred visual size of an object may arise as a consequence of perceptual processing (rather than explicit recognition and reasoning), where certain kinds of visual features are processed more effectively in certain visual sizes than other visual features. While it is still under active exploration what exactly these visual features existing in texform images are, it is nevertheless clear from the behavioral patterns discussed above that such features can drive real-world size effects at the level of perceptual processing. As such, it is possible that one of the underlying causes of the systematic canonical visual sizes are in fact perceptual in nature (which is not mutually exclusive with a role for knowledge in size preferences as well). Our goal in this study is to explore this possibility.

### 1.3 The current study: Perceptual contributions to canonical size?

Here, we tested if the canonical visual size of objects can be observed even when the images of objects have been “texformed”, so they are no longer recognizable, preventing explicit access to the objects’ identities and associated real-world size knowledge. We first asked in Experiment 1 if there are systematic canonical visual sizes for texform images, using a method of adjustment, which, to foreshadow, yielded equivocal results. We then turned to a forced-choice paradigm in Experiment 2a and its replication Experiment 2b, which showed clear and replicable results. In Experiment 3, we once again replicated the main results and ruled out alternative explanations.

## 2. Experiment 1: Method of Adjustment

In the first experiment, we examined whether intact and texform images have systematic canonical visual sizes using a method of adjustment task: Subjects were asked to rescale an image presented on the screen until it “looks best”. The key questions are: First, do we replicate Konkle and Oliva (2011), showing consistent preferred visual sizes for intact recognizable objects related to their real-world size? And, second, do texforms show consistent preferred visual sizes, related to real-world size, corresponding to the original images?

### 2.1 Method

#### 2.1.1 Participants

Fifteen naive subjects (6 females, 8 males, and 1 other gender; all with normal or corrected-to-normal visual acuity) from the Harvard University community completed individual 60-min sessions in exchange for a small monetary payment or a course credit. This sample size was preregistered**^1^** and was fixed to be identical across all experiments reported here. Four subjects were replaced based on predetermined exclusion criteria reported in Section 2.1.5 Exclusions.

#### 2.1.2 Apparatus

The experiment was conducted with custom software written in Python with the PsychoPy libraries (Peirce, Gray, Simpson, MacAskill, Höchenberger, Sogo, et al., 2019). The subjects sat approximately 60 cm without restraint from an iMac computer (with a viewport of 47.6 cm × 26.7 cm and effective resolution of 2048px × 1152px).

#### 2.1.3 Stimuli

The final stimulus set consisted of 40 original recognizable images and 40 corresponding texforms images (depicting 10 big animals, 10 big objects, 10 small animals, and 10 small objects). To generate this curated and controlled set of images, we used the following procedure.

First, a superset of 180 recognizable images were collected from various sources including stimuli from previous works (Long et al., 2018; Konkle & Caramazza, 2013; Konkle & Oliva, 2012) and Google images—consisting of 90 big items (big enough to support an adult human being) and 90 small items (small enough to be held by one hand), with an equal balance of animals and man-made objects. These images went through preprocessing to (a) remove the backgrounds, (b) crop to the smallest square that envelops the items, (c) resize to 512px × 512px, (d) convert to grayscale, (e) equalize their luminance and luminance histograms, and (f) place in the center of a gray background of 640px × 640px. The details of the preprocessing can be found in Appendix A. The resulting images are referred to as intact images (see Figure 1).

Next, the corresponding texform images (see Figure 1) were generated from these intact images following the method detailed in Deza et al. (2019), which is a variation and extension of the method used in Long et al. (2018). To overview, each texform image was synthesized from a random noise image seed, coerced to match the first and second order image statistics of each intact input image (following Freeman & Simoncelli, 2011). The size of the pooling windows over which these textural image statistics were computed reflects a peripheral placement in a simulated visual field (i.e., with small enough pooling windows with respect to the visual size of the depicted object to retain some coarse form information, but large enough with respect to the visual size of the depicted object to texturize the content, usually beyond recognition). Note this slightly modified texform algorithm enabled us to synthesize higher resolution texform images (640 px × 640 px) than in Long et al. (2018) (180 px × 180 px), for more on the method see Deza et al. (2019).

Finally, following the generation of these candidate texform images, we conducted an online pretest to test for texform recognizability (for details of the pretest, see Appendix B). Based on these results, we selected a final set of 40 pairs of intact and texform images (20 big items and 20 small items, half depicting animals and half depicting inanimate objects). The texform images were unrecognizable for at least 15 out of 18 pretest observers at the basic level (e.g., “dog” rather than “animal” or “huskie”; Rosch, Mervis, Gray, Johnson, & Boyes-Braem, 1976). This criterion was still a relatively low cut-off, so our experiment included a recognition post-test and excluded for each subject the images they recognized from the analysis.

#### 2.1.4 Procedure and Design

Each trial began with a 400 ms blank gray screen (matching the background color of all the images) followed by the presentation of a single centered image. The subjects were instructed to move the invisible cursor up and down to make size adjustments to the image (“adjust the size of the picture until it looks best to you…make the image the size you find most visually pleasing”). Moving the cursor up increased the visual size and moving it down decreased the visual size (see Figure 2a). The allowed size ranged from 5px × 5px to 1552px × 1552px. All images were initially presented at the medium size of 778px × 778px. The subjects made as many adjustments as they liked, to make the image the size they found “most visually pleasing”. They then clicked the mouse to submit their responses. (Mouse clicks within 300 ms of the onset of the images were recorded but ignored.)

**Figure 2.**
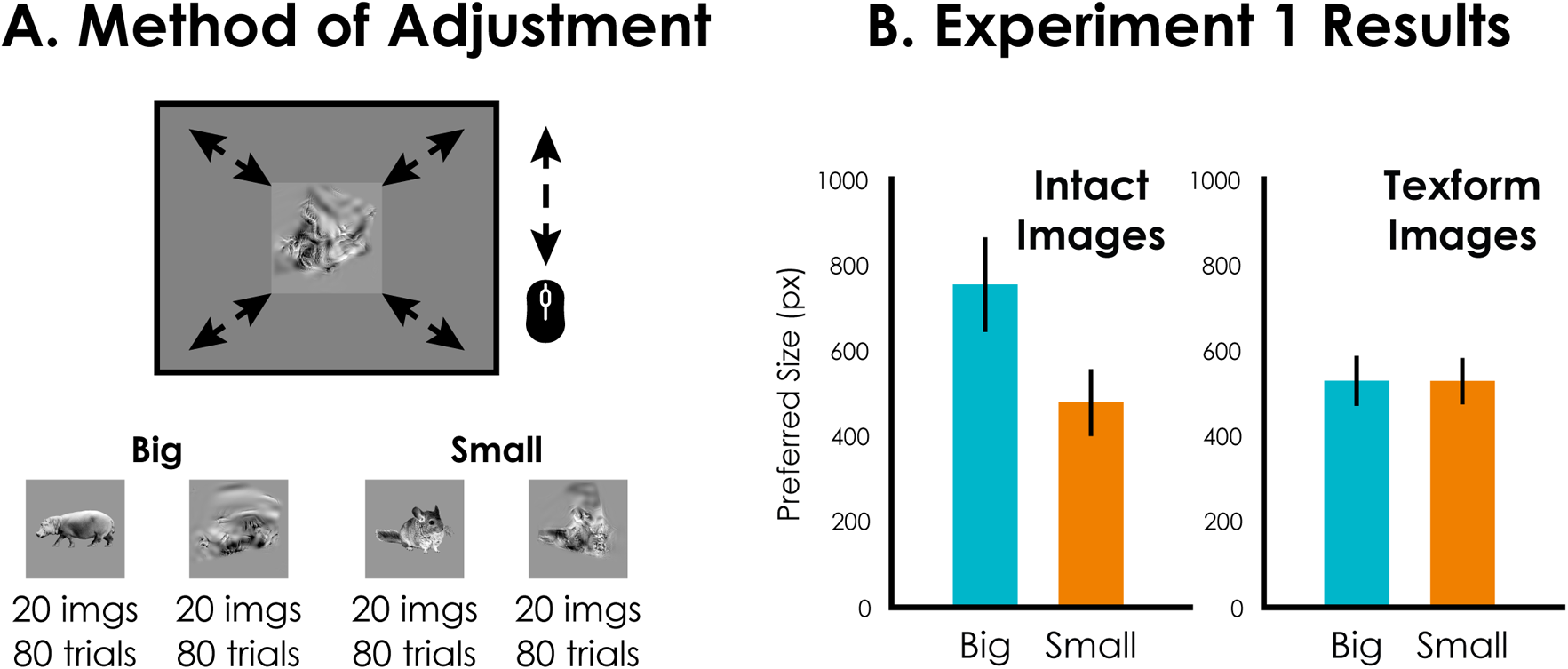
(a) Example displays of the size adjustment task in Experiment 1. (b) Results of the preferred visual size for big and small items, and intact (left) and texform images (right). Error bars reflect 95% confidence intervals for a within-subjects design.

In each “intact block”, all 40 intact images (2 real-world sizes × 20 items) were each presented once in a block in randomized order; and in each “texform block” the 40 texform images were presented in different randomized order. Five subjects completed four intact blocks followed by four texform blocks, and 10 subjects completed four texform blocks followed by four intact blocks. All completed a total of 320 trials. The block order alternated between subject before subject exclusions (data from all but one subject showed the same pattern in the critical analysis reported below, regardless of the block order). Subjects took three self-paced breaks when they completed 25%, 50%, and 75% of the experiment. Before the main experimental trials, subjects completed four practice trials with images (one big object, one big animal, one small object, one small animal, either all intact or all texform, depending on the first block type); these images never appeared in the main experimental trials. The subjects were not told about the nature of the texform images.

After the adjustment task, subjects completed a recognition test on all texforms to assess whether these specific subjects recognized the texform images (though note that these subjects also saw the corresponding intact images in the same setting). They were told that the texform images they saw were “made from images of objects by distorting the images while keeping their textures”. They then viewed all texform images one by one again and typed in with a keyboard what they thought was depicted in each image.

#### 2.1.5 Exclusions

The responses from the recognition test were graded by the first author before looking at the adjustment data: To be conservative at estimating the unrecognizability of texform images, any response that named an object with a similar size and shape from the depicted object was considered correct. Any adjustment trials from stimuli with their texform version recognized in the recognition test were discarded, along with adjustment trials with mouse clicks within 300 ms of the onset of the images. Three subjects had more than 20% of trials discarded and thus were excluded and replaced with new subjects to meet the targeted sample size.

Next, we tested the consistency of the preferred visual sizes of the intact images since this is a necessary precondition for examining textform feature contributions to this preferred visual size. To estimate the reliability of the preference, we computed the correlation between the selected sizes across the first half and the last half of trials for the intact, recognizable images. One subject was removed based on having low reliability (*r’*<.5), and was replaced, yielding a final average reliability of the preferred visual sizes for intact images of *r’*=.82 (*SD*=.15). Unlike the preregistered plan, we did not exclude subjects based on the reliability of the preferred visual size of texforms because the reliabilities were generally very low (*r’*=.21, *SD*=.25). After subject replacements, the mean recognition rate was 7.3% (*SD*=5.9%) and a total of 68 out of 4800 trials were discarded due to early mouse clicks.

### 2.2 Results and discussion

First, we examined whether, for intact recognizable objects, subjects consistently preferred visual sizes that were related to the real-world size of the depicted object. The results are shown in Figure 2 (individual data for all experiments are shown in Appendix C). Overall, we found that people did show the signature preferences. Subjects preferred to view the big items at a bigger visual size (757px, *SD*=268px) compared to small items (480px, *SD*=180px; *t*(14)=3.78, *p*=.002, Cohen’s *d*=0.98; 14 out of 15 subjects, *p*=.002), regardless of whether the images depicted animals or inanimate objects (main effect of Animacy: *F*(1,14)=0.94, *p*=.348, *η_p_*^2^=.063; interaction effect between Animacy and Size: *F*(1,14)=3.45, *p*=.085, *η_p_*^2^=.198). Thus, these results are consistent with the canonical visual size effect (Konkle & Oliva, 2011).

On the other hand, with the texform images, the reliability of the preferred sizes was quite low (*r’*=.21, *SD*=.25), indicating subjects did not select similar visual sizes across repeated presentations of the same texform image. Further, we did not observe the signature preference where the big items were preferred at bigger visual sizes than small items (531 px, *SD*=153, vs 530 px, *SD*=160; *t*(14)=0.07, *p*=.944, Cohen’s *d*=0.02; 10 out of 15 subjects, *p*=.302). The 2 (big/small) × 2 (intact/texform) ANOVA showed a significant main effect of size (*F*(1,14)=14.92, *p*=.002, *η_p_*^2^ =.516), no main effect of intact/texform (*F*(1,14)=2.9, *p*=.110, *η_p_*^2^=.172), and a significant interaction effect (*F*(1,14)=13.1, *p*=.003, *η_p_^2^*=.483). Thus, the simple act of resizing until the texform image “looks best” did not yield consistent preferred visual sizes.

While these results could indicate an actual lack of size preference for texform images, the unreliable responses may also be related to the nature of the adjustment task. For example, in facing the unfamiliar texform images, it is possible that subjects felt less confident in making a choice from the unlimited options given by an adjustment task, leading to a family of unconstrained strategies. We thus performed Experiment 2a and 2b with a more rigorous psychophysical method to probe for the existence of visual size preferences in texform images. Additionally, the lack of effect in this adjustment task has one interpretive benefit—that is, it provides further support that the subjects are not systematically recognizing these texform images as something (if they were, the sizes would be consistent across repetitions).

## 3. Experiment 2: Forced-Choice Task

Experiment 2a and 2b cut the number of preferred visual size options down from unlimited to only two, using a forced choice paradigm. That is, subjects could toggle between two options and selected the one that looked best. We conducted two versions of this experiment: In the first version, we created a larger stimulus set drawn from the same superset as reported in the experiment above. In the second version, intended as a replication experiment with some generalization, we changed the stimulus set again, in order to dovetail more closely with previous work, using a subset of the original texform images used by others (Long et al., 2016; Long & Konkle, 2017; Long et al., 2018; Wang, Janini, & Konkle, 2022; Grootswagers, Robinson, Shatek, & Carlson, 2019; see Figure 1). These two versions of the experiment (Experiment 2a and 2b) were otherwise identical, except for the stimuli used and a few related details in their presentation.

### 3.1 Method

The experimental apparatus and general procedures were similar to Experiment 1, except as noted here.

#### 3.1.1 Participants

Each experiment was completed by 15 naive subjects. (Experiment 2a: 10 females, 5 males; Experiment 2b: 6 females, 9 males). Subjects were replaced based on preregistered**^2^** exclusion criteria reported in Section 3.1.4 Exclusions (6 excluded and replaced in Experiment 2a; 1 excluded and replaced in Experiment 2b). All experiments were approved by either the Harvard University or the UCLA Institutional Review Board.

#### 3.1.2 Stimuli

In Experiment 2a, 50 pairs of intact and texform images were repicked from the superset of 180 processed images as described in Experiment 1. Half of the images depicted big items and half small items (with a balanced selection of animals and inanimate objects). Based on a pilot study using the same adjustment task from Experiment 1, these images were selected to maximize the range of canonical visual sizes, while also maintaining a generally balanced set across real-world size and animacy (12 big and 13 small animals; 12 big and 13 small objects). The images were then scaled down to 440px × 440px, which is a lower resolution than in Experiment 1 (based on pilot studies, this design choice helped to ensure that the preferred sizes of intact images were well within the size of the screen).

In Experiment 2b, 50 pairs of images were selected from the stimuli from Long, et al. (2018), available online (https://konklab.fas.harvard.edu/). The main difference of these texforms is that they have a lower spatial resolution.**^3^** The images were selected to include 25 animals and 25 objects and maximize the range of canonical visual sizes based on a pretest, this resulted in 19 big items (8 big animals, 11 big objects) and 31 small items (17 small animals, and 14 small objects). Note that, here the division of items into “big” and “small” is less relevant, as we can treat size here as a continuous variable.

#### 3.1.3 Procedure and Design

The subjects completed three tasks in order: (a) an adjustment task on intact images, to obtain a canonical visual size estimate for each item, (b) the main forced choice task, to select which of two visual sizes of the same image was more aesthetically pleasing, completed for both intact and texform images in different blocks, and (c) a post-test assessing recognition on the texform images. This procedure is depicted in Figure 3a.

**Figure 3.**
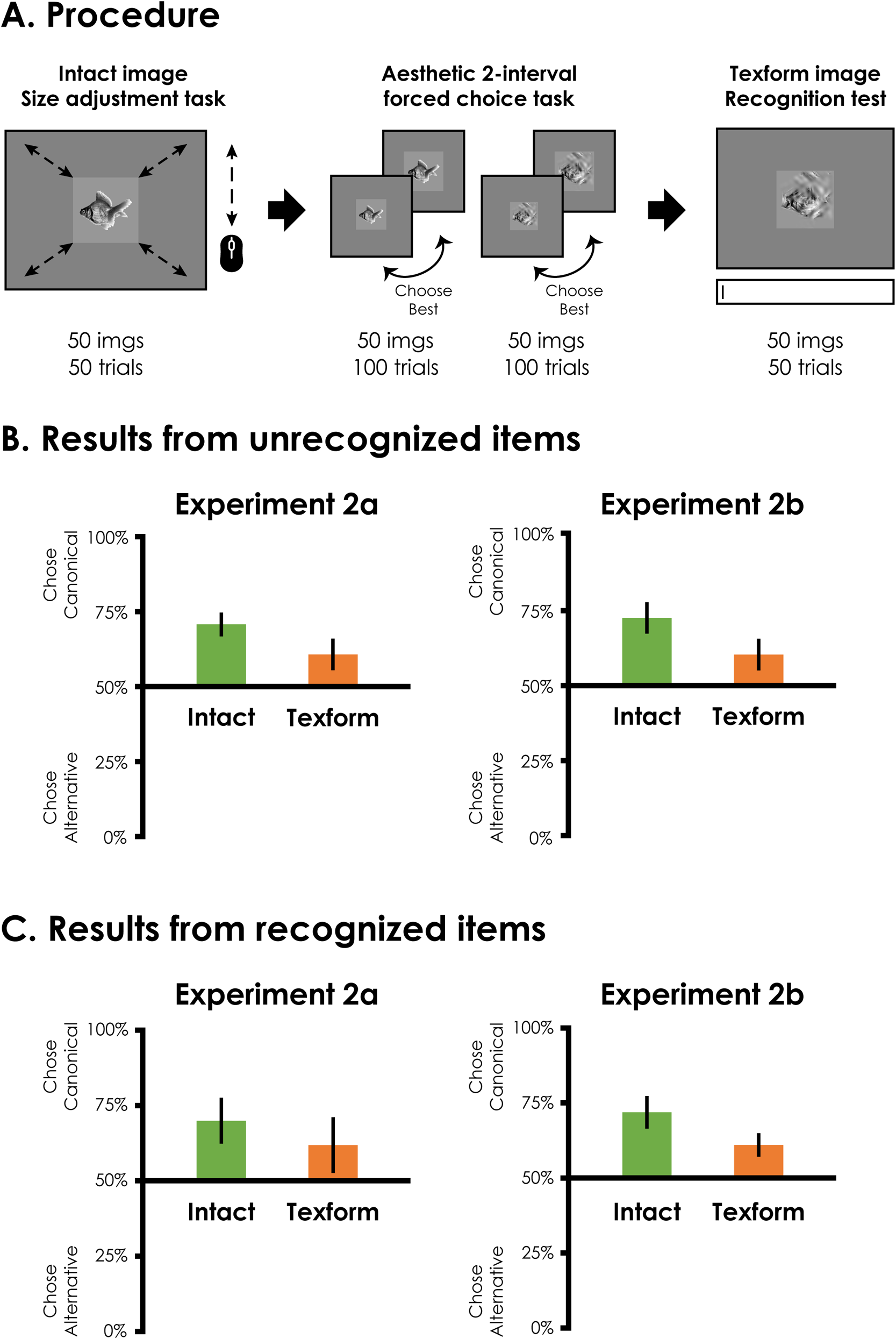
(a) Subjects completed 3 tasks in order in Experiment 2a and 2b: Size adjustment task on intact images, followed by an aesthetic 2-interval forced choice task, followed by a texform image recognition test. (b) Visual size preferences for the set of images whose texforms were not subsequently recognized, for Experiment 2a (left) and 2b (right). The y-axis shows the percentage of trials the subjects chose the canonical visual size (chance = 50%), plotted separately for intact and texform images. Error bars reflect 95% confidence intervals. (c) The plots are the same as in (b), but for the subset of items for which the texforms were subsequently recognized.

##### Size adjustment task on intact images

As in Experiment 1, subjects moved the mouse to adjust the visual size of an image on the screen and clicked when the image “looked best.” This task was identical to the adjustment task in Experiment 1, except that it consisted of only four intact blocks of 50 trials, the allowed size ranged from 20px × 20px to 2152px × 2152px, with images initially presented in 20px × 20px (Experiment 2a) or 1086px × 1086px (Experiment 2b), the click detection started earlier at 100 ms after the onset of the images, and there was only one practice trial for Experiment 2b. For each item, its canonical visual size for that subject was calculated by averaging the selected sizes from the repetitions, after excluding trials with a response time (RT) less than 300 ms and excluding items that had more than half of the four trials excluded. Only the items that yielded canonical visual size smaller than 871px × 871px (so that the images stayed well within the monitor’s size) entered the next task (with both the intact images and their corresponding texform images). Critically, these canonical visual sizes were used to set the choice options for the next task.

##### Aesthetic two-interval forced choice task

On each trial, subjects viewed a single image and toggled between two sizes with a key press. They were asked to toggle to view both sizes as many times as they liked and decide which of the sizes “looks more aesthetically pleasing”. Unbeknownst to the subjects, one of the size options was the average canonical visual size they picked for that item in the adjustment task, and the other was a 30% difference in the diagonal length (with each item presented with a visually bigger alternative option in one trial, and visually smaller alternative option in another trial; see Figure 3a for example displays from Experiment 2a and 2b). Which size option was shown first in a trial was randomized.

Subjects performed this task in separate blocks for intact and texform images. Critically, in the texform block, the visual sizes were based on the canonical visual sizes of the corresponding intact images. For both intact and texform blocks, we calculated the percentage of trials in which subjects picked the canonical visual size options as the key outcome measure. The block order alternated between subjects before subject exclusions (resulting in nine subjects completing the texform block first, and six completing intact block first in Experiment 2a; with seven completing the texform block first and eight completing the intact block first in Experiment 2b).

##### Texform image recognition test

Finally, as subjects received extensive exposure to the intact images as well as the corresponding texforms, we next tested the recognizability of the texforms that were included in the forced choice task. The same procedure and grading as in Experiment 1 were performed, and the average texform image recognition rate was 24% (*SD*=14%) for Experiment 2a and 56% (*SD*=12%) for Experiment 2b. This difference in recognition rates was inherent to the different stimulus sets, and did not lead to qualitative differences in the results. Below we report analyses from unrecognized and recognized items separately.

#### 3.1.4 Exclusions

The following preregistered exclusion criteria were applied: (a) forced choice trials with RT less than 300 ms, (b) forced-choice trials without any toggling (i.e., the subject picked the first option without viewing the second option), (c) subjects with more than or equal to 5% trials excluded in either the adjustment or the forced choice task (Experiment 1a N=1; Experiment 1b N=1), (d) subjects who had less than 12 items (i.e., 12 intact and 12 texform images)**^4^** entered into the forced choice task (Experiment 1a N=1), and (e) subjects who had by-item split-half reliability lower than 0.5 in the adjustment task (Experiment 1a N=4). As noted above, excluded subjects were replaced to achieve the pre-registered N=15 for each experiment.

#### 3.1.5 Analysis

Our analysis plan followed this rationale: If explicit (non-perceptual) size knowledge is required for viewing size preference to arise, then we should not see similar canonical viewing sizes for intact images and unrecognized texform images. Thus, we first considered only the forced-choice trials that the texforms were not subsequently recognized during the recognition test, and calculated the percentage of trials in which subjects chose the canonical visual size rather than the modulated size. Then, we simply compared these percentages with the chance level of 50% with a one-sample t-test. We did this analysis separately for the intact images and the texform images.

For exploratory purposes, we also performed the same analyses on the recognized texform images for the readers’ information. These exploratory analyses were not part of our preregistered analysis plan, and our experiment was not designed for them: We have specifically generated texform images to be relatively unrecognizable for our main analyses, which necessarily led to low numbers of recognizable texform images. We nevertheless reported these analyses in case they may inform future studies. None of our major conclusions depend on the results of these analyses.

### 3.2 Results and discussion

#### 3.2.1 Main analyses with unrecognized items

We first analyzed the visual preference data from the forced choice task, but only including trials in which the texforms were not subsequently recognized during the recognition test. (This included an average of 76.9% of the forced-choice trials in Experiment 2a, 43.9% in Experiment 2b.) The percent of trials in which subjects chose the canonical visual size rather than modulated size was plotted for both intact and texform blocks in Figure 3b.

Inspection of the figure reveals two main results, present in both experiments. First, subjects chose the canonical size over the alternative above chance, for intact images and, critically, also for texform images. Second, the canonical visual size preference was stronger in intact than in texform images. Indeed, one-sample t-tests confirmed all blocks were above the chance level of 50% (Experiment 2a: intact block, 71%, *SD*=7%, *t*(14)=10.90, *p*<.001, Cohen’s *d*=2.81, 15 out of 15 subjects, *p*<.001; texform block, 61%, *SD*=9%, *t*(14)=4.38, *p*=.001, Cohen’s *d*=1.12, 13 out of 15 subjects, *p*=.007; Experiment 2b: intact block, 73%, *SD*=9%, *t*(14)=9.24, *p*<.001, Cohen’s *d*=2.38, 14 out of 15 subjects, *p*=.001; texform block, 60%, *SD*=10%, *t*(14)=4.19, *p*=.001, Cohen’s *d*=1.08, 12 out of 15 subjects, *p*=.035), and the differences between intact and texform blocks were also significant (Experiment 2a: *t*(14)=3.16, *p*=.007, Cohen’s *d*=0.82, 13 out of 15 subjects, *p*=.007; Experiment 2b: *t*(14)=3.81, *p*=.002, Cohen’s *d*=0.98, 12 out of 15 subjects, *p*=.035).**^5^** We also conducted a 2×2 ANOVA (with Experiment 2a/2b as a between subject factor and texform/intact block as a within-subject factor), which was not pre-registered, to provide additional information for interested readers. We found a significant main effect of block type (*F*(1,27)=24.2, *p*<.001, *η_p_*^2^=.472), no main effect of experiment (*F*(1,27)=0.9, *p*=.339, *η_p_*^2^=.034), and no interaction effect (*F*(1,27)=0.1, *p*=.756, *η_p_^2^*=.004).

These findings confirmed that subjects preferred to view texform images in the canonical visual size of their original versions compared to bigger or smaller alternatives. Thus, visual size preferences partially persist after object recognition is disrupted, providing evidence that mid-level visual features contribute to the phenomenon of canonical visual size. However, at the same time, the visual size preference was still stronger in intact images. What may explain this difference? One possibility is that the difference still stemmed from the visual features: While texform images were made to be unrecognizable but preserve visual features, it is impossible to preserve all visual features. Among these visual features that are only available in the intact images, some may contribute to the viewing size preference through perceptual routes. Another possibility is that the recognition and thus access to object knowledge strengthen the viewing size preference in intact images. The following exploratory analyses with recognized items help shed light on these possibilities.

#### 3.2.2 Exploratory analyses with recognized items

The same analyses were performed on trials with recognized items (see Figure 3c). Inspection of the figure suggested the same three trends in both experiments: First, subjects chose canonical size over the alternative above chance level with both intact and texform images. Second, canonical size preference was stronger in intact than in texform images. Third, the data patterns were almost numerically identical to the unrecognized trials. One-sample t-tests confirmed the first impression that all blocks was above the chance level of 50% (Experiment 2a: intact block, 70%, *SD*=14%, *t*(14)=5.63, *p*<.001, Cohen’s *d*=1.45, 12 out of 15 subjects, *p*=.035; texform block, 62%, *SD*=17%, *t*(14)=2.75, *p*=.016, Cohen’s *d*=0.71, 10 out of 15 subjects, *p*=.302; Experiment 2b: intact block, 72%, *SD*=10%, *t*(14)=8.60, *p*<.001, Cohen’s *d*=2.22, 15 out of 15 subjects, *p*<.001; texform block, 61%, *SD*=7%, *t*(14)=5.99, *p*<.001, Cohen’s *d*=1.55, 13 out of 15 subjects, *p*=.007). There was also a difference between intact and texform blocks, but this difference was only significant in Experiment 2b (Experiment 2a: *t*(14)=1.37, *p*=.193, Cohen’s *d*=0.35, 8 out of 15 subjects, *p*>.999; Experiment 2b: *t*(14)=3.07, *p*=.008, Cohen’s *d*=0.79, 11 out of 15 subjects, *p*=.118). This is likely due to the lower recognition rate in Experiment 2a than in 2b (24% vs. 56%), leading to fewer trials entering the analysis and thus higher variances (*SD_diff_*=23% vs. *SD_diff_*=12%). With another 2×2 ANOVA with experiment and texform/intact block types as factors, we found a significant main effect of block type (*F*(1,27)=6.5, *p*=.016, qa*η_p_*^2^=.195), no main effect of experiment (*F*(1,27)=0.4, *p*=.526, *η_p_*^2^ =.015), and no interaction effect (*F*(1,27)<0.1, *p*=.831, *η_p_^2^*=.002).

While remembering to take cautions on the tentative nature of these observations, the almost identical numerical patterns found in unrecognized and recognized images suggests that explicit identity and/or size knowledge access may not be a major factor in the effects found here. Texform images still showed weaker canonical visual size preferences than intact objects, even when the texforms were subsequently recognized (and thus could have potentially allowed access to physical size knowledge during the aesthetic choice task). These results provide additional support that these consistent visual size preferences for texform images are driven by their visual features.

## 4. Experiment 3: Priming Control

While the results from the last experiments were encouraging, there remains an alternative explanation to the above-chance performances on the texform choice task: It is possible that when subjects were selecting which of the two sized-texform views looked best, the tendency to pick the canonical view that matched with the intact item was induced by priming. That is, perhaps participants were remembering (either implicitly or explicitly) the pairing of adjusted sizes and image features when selecting the preferred views of intact images, in turn priming their choices in the texform choice task. Thus, to understand the degree to which priming could explain these results, here we designed a task to directly rely on priming to explore whether this mechanism could account for the consistent texform viewing preferences.

Subjects participated in one of two task conditions, in a between-subjects design: The aesthetic task and the priming task. The aesthetic task was a replication of Experiment 2a, while the priming task measured the priming effect directly by not asking for aesthetic judgments but instead asking subjects to pick the sizes that were paired with the images in the adjustment task. We compared the two task conditions to see if a priming effect can explain the canonical viewing size preferences in texform images observed above. Additionally, we also only presented texform images during the visual choice blocks (removing intact images in this phase of the experiment) to ensure that those extra exposures to the intact images were not critical to our findings.

### 4.1 Method

The experimental apparatus and general procedures were identical to Experiment 2a, except as noted.

#### 4.1.1 Participants

Each task condition was completed by 15 naive UCLA undergraduate students (all 30 subjects happened to be female). Subjects were replaced based on exclusion criteria reported in Section 4.1.4 Exclusions (three subjects in the aesthetic task condition and three subjects in the priming task condition were excluded and replaced).

#### 4.1.2 Apparatus

The subjects sat approximately 60 cm without restraint from a Dell computer (with a viewport of 41.0 cm × 31.0 cm and effective resolution of 1600px × 1200px.

#### 4.1.3 Procedure and Design

##### Aesthetic task condition

The same three phases from Experiment 2a were performed with slight adjustments to accommodate the new apparatus: For the size adjustment task on intact images, the allowed size ranged from 20px × 20px to 2200px × 2200px, with images initially presented in 20px × 20px. Only the items that yielded canonical visual size smaller than 908px × 908px (so that the images stayed well within the monitor’s size) entered the next task. For the two-interval forced choice task, subjects only performed this task on texform images. Each texform image were tested four times, with two trials showing the 30% bigger alternative and two trials showing the 30% smaller alternative, with order of size option in two intervals counterbalanced. For the texform image recognition test, the average recognition rate was 16% (SD=10%).

##### Priming task condition

The three phases of the experiment were similar to the aesthetic task condition with the following critical differences. First, during the size adjustment task on intact images, subjects were asked to adjust the image to the size of a box (10px red border) on the screen (rather than the best-looking size; “adjust the size of the picture until the picture fit right into the square…adjust the picture to the largest size that still fits in the box”). The sizes of the box for each image were yoked to the canonical size selected by subjects in the aesthetic task condition. For example, we took the canonical sizes picked in the adjustment task by Subject #1 in the aesthetic task condition, randomized the pairing between canonical sizes and the images, and had Subject #1 in the priming task condition adjust the images to the new paired sizes (indicated by the box size). The motivation here is to allow participants to (explicitly or implicitly) learn an association between an intact item and a visual size that is free of aesthetic biases. This way, the subsequent choice task will reflect the strength of priming effect rather than any aesthetic preferences.

There were four practice trials, with the box size set to the median of all sizes in the formal trials. Because there was no way to show a box larger than the screen, for canonical sizes bigger than the size limit imposed in the aesthetic task condition, the box was presented in the average size of all sizes instead. (Note that these trials were merely fillers to equate the subjects’ experience between the two conditions. These stimuli were not included in the two-interval forced-choice task as in the aesthetic task condition.) Only when the subjects adjusted the image to the correct size (within 5% error margin), the box turned green and mouse clicks were allowed for submitting the response.

Next, for the two-interval forced choice task, subjects were asked to view the texforms in two different sizes, and chose the size that their intact counterparts were adjusted to in the adjustment task (“Your job is to look at both of the displays of the same picture in different sizes, and choose the size that corresponds to the size its undistorted version was adjusted to in the first part”). Each subject was tested on texforms retained for the corresponding subject in the aesthetic task condition.

Finally, participants completed the same texform image recognition test, the average recognition rate was 19% (*SD*=14%).

#### 4.1.4 Exclusions

The following exclusion criteria were applied: (a) forced choice trials with RT less than 300 ms, (b) forced-choice trial without any toggling (i.e., the subject picked the first option without viewing the second option), (c) subjects with more than or equal to 5% trials excluded in either the adjustment or the forced choice task (Priming N=2), (d) subjects who had less than 12 items entered into the forced choice task (Aesthetics N=3), and (e) an experimenter error resulting in running a repeating subject (Priming N=1). As noted above, excluded subjects were replaced to achieve N=15 for each task condition.

#### 4.1.5 Analysis

The same main and exploratory analyses from Experiment 2a and 2b were applied to both the aesthetic and priming task conditions, except for two differences: First, there were no intact images tested here. Second, the t-tests used to compare the results from two task conditions were two-sample t-tests because of the between-subject design. Again, none of our conclusions rely on the results of the exploratory analyses.

### 4.2 Results and discussion

#### 4.2.1 Main analyses with unrecognized items

We first analyzed the visual preference data from the forced choice task, but only including trials in which the texforms were not subsequently recognized during the recognition test. (This included 84.4% of the forced-choice trials in the aesthetic task condition, 81.4% in the priming task condition.) The percent of trials in which subjects chose the canonical visual size rather than alternative size was plotted in Figure 4.

**Figure 4.**
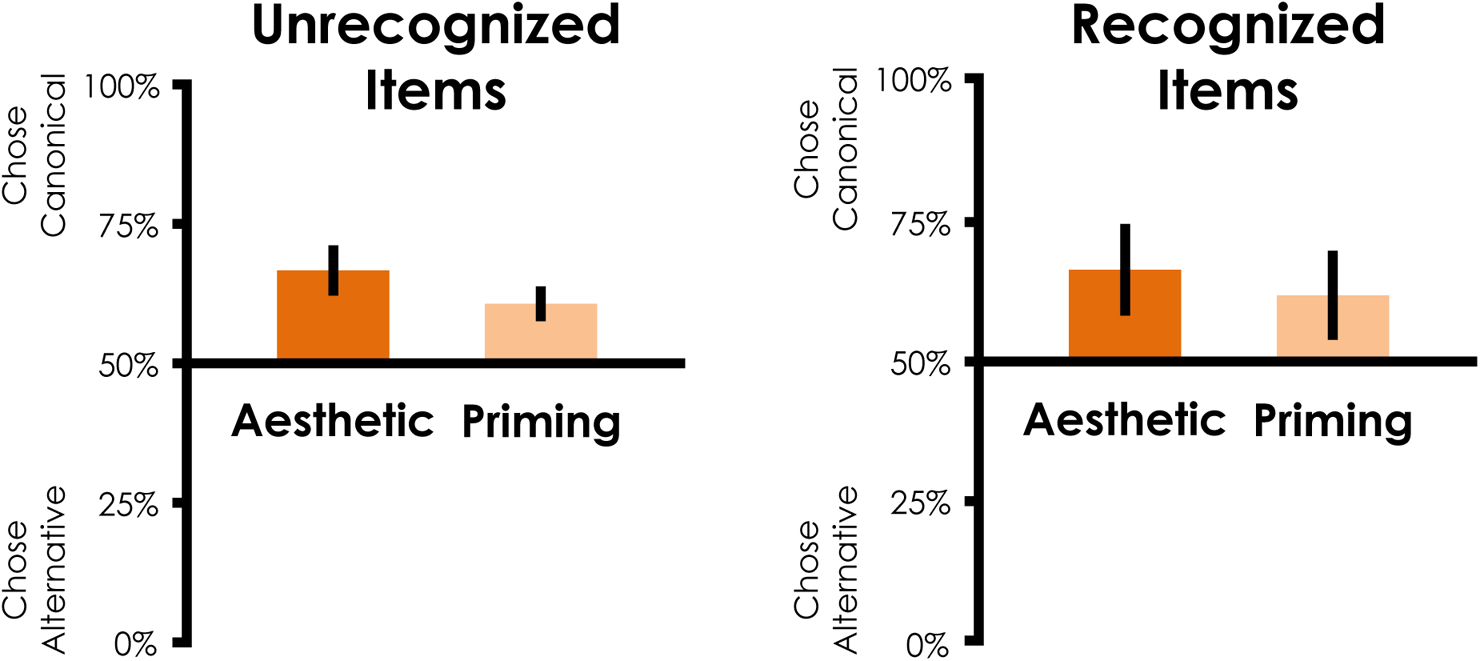
Visual size preferences for unrecognized (left) and recognized (right) items for Experiment 3. The y-axis shows the percentage of trials the subjects chose the canonical visual size (chance = 50%), plotted separately for the group of subjects competing the aesthetic task condition, and the group of subjects completing the priming task condition. Error bars reflect 95% confidence intervals.

Inspection of the figure reveals two patterns: First, subjects chose the canonical size over the alternative above chance in both the aesthetic and priming task conditions. Second, subjects in the aesthetic task condition picked the canonical sizes more readily than subjects in the priming task condition. One-sample t-tests confirmed the first impression: Both conditions were above the chance level of 50% (Aesthetic: 66.7%, *SD*=8.1%, *t*(14)=7.97, *p*<.001, Cohen’s *d*=2.06, 15 out of 15 subjects, *p*<.001; Priming: 60.7%, *SD*=5.6%, *t*(14)=7.38, *p*<.001, Cohen’s *d*=1.90, 15 out of 15 subjects, *p*<.001). Critically, the difference between conditions were significant (*t*(28)=2.35, *p*=.026, Cohen’s *d*=0.86). These findings confirmed that (a) subjects preferred to view texform images in the canonical visual size of their original versions compared to bigger or smaller alternatives, replicating Experiment 2a and 2b (observed from the aesthetic task condition), (b) part of this effect can be attributed to a priming effect (demonstrated in the priming task condition), but most importantly, (c) the priming effect did not fully explain the effect in the aesthetic task condition. Thus, over and beyond a priming effect from size pairing exposure in the adjustment task, visual size preference persists after object recognition is disrupted. This strengthened the evidence that mid-level visual features contribute to the phenomenon of canonical visual size.

#### 4.2.2 Exploratory analyses with recognized items

The same exploratory analyses were performed on trials with recognized items (see Figure 4. Inspection of the figure suggested a clear pattern: Subjects chose canonical size over the alternative above the chance level (Aesthetic: 66.3%, *SD*=14.1%, *t*(13)=4.32, *p*=.001, Cohen’s *d*=1.16, 12 out of 14 subjects, *p*=.013, one subject did not recognize any texform; Priming: 61.8%, *SD*=14.3%, *t*(14)=3.19, *p*=.007, Cohen’s *d*=0.82, 9 out of 15 subjects, *p*=.607). However, there were no difference between the two conditions (*t*(27)=0.86, *p*=.398, Cohen’s *d*=0.32), which is likely due to the low recognition rate and thus high variances (*SD_pooled_*=14.2%). Again, the results from unrecognized and recognized items were very similar (Aesthetics: *M_Diff_*=0.6% (13.8%), *t*(13)=0.16, *p*=.876, Cohen’s *d*=0.04, excluding the subject who recognized none of the texforms; Priming: *M_Diff_*=1.0% (15.7%), *t*(14)=0.26, *p*=.801, Cohen’s *d*=0.07), suggesting that explicit identity and/or size knowledge access is not a major factor in the effects found here.

## 5. General Discussion

Our minds have at least two sources of information when it comes to representing the physical size of objects in the world: We can access knowledge about the objects’ size attributes from knowing what they are (e.g., Chen et al., 2014), and we also perceive visual feature differences between objects of different sizes (e.g., Long et al., 2016). Here, we asked what kind of information is driving the systematic visual size preference, where we like to view big things big and small things small (Konkle & Oliva, 2011; Linsen et al., 2011). In three experiments, we first replicated the systematic visual size preferences for recognizable objects and found some evidence for the role of perceptual features in such preferences: While resizing texforms until they looked best did not show visual size preferences (Exp 1), we did find consistent preferences when given only two options (replicated across Exp 2a and 2b), which could not be fully explained away by priming mechanisms between intact and texform images (Exp 3). These results demonstrate that visual size preferences, instead of only stemming from knowledge of objects’ physical sizes, can also be evoked by the visual features that are preserved in unrecognizable texforms.

### 5.1 Mid-level visual features: What are they?

Since the visual size preferences for texforms are systematic without object recognition, the information about the objects’ physical sizes must be coming from the visual features. What kind of visual features carry the information about an object’s physical size? While our experiments do not have direct evidence to pinpoint the responsible features, the use of texform images constrained the possibilities. First, the effects cannot be explained by color or global low-level visual statistics (e.g., luminance and contrast), since the images were converted to grayscale and equalized across luminance and luminance histograms (see Appendix A). Second, the effects do not rely on accurate identification of the texform images, and thus cannot be attributed to explicit semantic information regarding the particular objects. These bounds leave a wide range of “mid-level” features in between. In our case, texform images would presumably be on the lower-end of this range, as these were created by matching the first- and second-order statistics of intact images within a series of receptive field-like pooling windows (Freeman & Simoncelli, 2011; Long et al., 2016), the global forms (rough spatial envelopes for the whole objects and their parts) are preserved along with local corners, junctions, and contours, but clear outer contours and three-dimensionality are less preserved.

One proposed mid-level visual feature dimension, which co-varies with the real-world size of objects, is related to perceived curvature (Konkle, 2011; Long et al., 2016). For example, on a 5-point likert scale from “very curvy” to “very boxy”, people consistently judged small objects to be curvier than big objects in both intact and texform images (Long et al., 2016). This subjective curvature dimension also predicts the structure of ventral visual system’s responses to objects that vary in real-world sizes (Long et al., 2018; see also Srihasam, Vincent, & Livingstone, 2014; Yue, Robert, & Ungerleider, 2020). Curvature computations more generally have been proposed to emerge from even earlier computations linked to spatial frequency and end-stopping that vary from the center to the periphery (e.g., Ponce, Hartmann, Livingstone, 2017). However, note that even “curvature” itself is a multi-level construct with a more primitive perceptual instantiation based on constructing adjacent orientations in a retinotopic format (e.g., Yue et al., 2020), to a more 3-dimensional representation of curvature in an object-centered format (e.g., Srinath, Emonds, Wang, Lempel, Dunn-Weiss, Connor, & Nielsen, 2021). Understanding these mid-level features, both visualizing them and developing a vocabulary to describe them, is still an active front of research, with potentially promising new in-roads through an analysis of the feature tuning across different layers of deep neural networks (e.g., Olah, Mordvintsev, & Schubert, 2017; Bau, Zhu, Strobelt, Lapedriza, Zhou, & Torralba, 2020)

Finally, there is an intriguing link between curvature and overall aesthetic experience, where curvier things generally give rise to a relatively positive aesthetic experiences (Bar & Neta, 2006; Cotter, Silvia, Bertamini, Palumbo, & Vartanian, 2017; Palumbo, Rampone, Bertamini, Sinico, Clarke, & Vartanian, 2020; Vartanian, et al., 2013; but see also, Maezawa, Tanda, & Kawahara, 2020).**^6^** Here we are speculating that feature variation along a dimension from curvy-to-boxy is systematically linked to preferences for smaller-to-larger visual sizes. Exactly how curviness and general aesthetic experience are related to visual size preferences remains an open empirical question.

### 5.2 Size knowledge’s role in aesthetics?

While we showed that pure perceptual processes contribute to canonical visual size preferences, we do not mean to imply that these solely determine canonical visual sizes. It is likely that knowledge of physical sizes still plays a (potentially substantial) role in size preference in other contexts. In fact, telling people that objects they were viewing were “toys” (thus were physically small) reduced the canonical visual sizes by more than 50% (Konkle & Oliva, 2011, Experiment 4). Our results from Experiment 1 may also imply effects of recognition and knowledge access on visual size judgements, as subjects showed clear reliable viewing size preferences with intact images. (However, note that it is theoretically possible that these visual size preferences for intact objects observed in an adjustment task are still related to underlying perceptual differences, just those that are not retained in texform images). Size knowledge is also known to influence aesthetic experience in a very different way—through expectations and pleasant surprises. Famously, artists (e.g., Claes Oldenburg) created humongous statues of everyday objects, which induced aesthetic experience presumably through challenging our expectations (e.g., Van de Cruys & Wagemans, 2011). These kinds of aesthetic experiences have been argued to differ in intensity (and maybe in nature as well) from those that we may rely on to pick out a canonical visual size that simply “looks good” (e.g., for a discussion on these different kinds of aesthetic experiences, see Makin, 2017; Brielmann & Pelli, 2017).

Thus, we are not trying to imply that recognition and explicit knowledge of identity or real-world size information have no role in shaping visual size preferences. Instead, we aim to remove those factors, and show that they do not fully account for visual size preferences, revealing evidence for the role of perceptual mechanisms as well.

### 5.3 The function of canonical visual sizes?

Why do we have canonical visual sizes? Of course, our study does not provide a direct answer, but inspires some speculative ideas. One possibility is that it is simply a byproduct of ontogenetic and/or phylogenetic developments of visual systems: The visual features’ correlation with physical sizes gives rise to correlation with visual sizes in experience as well. For example, if we tend to see small objects in smaller visual sizes, and small objects tend to be curvier, our visual systems may process visually small curvatures more fluently than visually big curvatures. And this perceptual fluency leads to a more positive experience when viewing physically small objects in small visual sizes (as in Reber, Schwarz, & Winkielman, 2004). In this way, the size preference itself may not have a particular function but is just an indication of the visual system’s tuning for features it commonly encounters.

Another possibility is that canonical visual size is in fact functional. For example, it may guide us to seek some sort of optimal viewing distances for each object (cf. Merleau-Ponty, 1962). A functional viewing distance might be one that minimizes the danger associated with getting close to unknown objects in the environment, yet close enough to gather information for appropriate actions. For example, if we are looking at an unknown object, our knowledge cannot guide us to interact with it in a proper distance, but our visual systems may use heuristics based on visual features to induce aesthetic experience, which in turns motivate us to seek a proper viewing distance.

Yet another possibility is that a functional viewing distance might be one that is linked to spatially varying sensitivity across the visual field. Being too far from an object prevents us from discerning important detailed features at sufficient resolution (e.g., different patterns on a leaf can help identify a poisonous plant). Being too close prevents us from seeing the global contours and summary statistics (e.g., how abundant a fruit tree is). The geometric relationship between visual size and viewing distance determines the proper viewing distance on this account, where smaller objects require a closer distance, and bigger objects demand a farther distance to project to appropriate visual sizes in the visual field**^7^**. More generally, a functional account argues that visual size preferences are there to assist active learning by motivating us to modulate the visual inputs themselves, adding support to the idea that aesthetic experience interacts with perception (e.g., Chen & Scholl, 2014; Chen, Colombatto, & Scholl, 2018; Forman, Chen, Scholl, & Alvarez, 2021;) and serves adaptive functions (e.g., Bar & Neta, 2006; Orians & Heerwagen, 1992).

The idea that aesthetic preferences are evolved to guide our action-perception cycle is closely related to a flavor of the predictive processing theory for affective values (e.g., Van de Cruys, 2017). This theory posits that positive aesthetic experiences arise from reducing either long-term or short-term prediction errors through information gain. Thus, viewing an object in the canonical viewing distance and canonical viewing size can maximize the information gain and subsequently reduce prediction errors, leading to positive experiences (for how this may work in a different phenomenon, see Van de Cruys, Damiano, Boddez, Król, Goetschalckx, & Wagemans, 2021). There are important bridges to be formed between these ideas of active sensing, optimal sensing, and aesthetic preferences: For example, understanding the degree to which predictive processes are key to the formation of canonical visual sizes, and the degree to which these mechanisms operate over pure perceptual representation or require the semantic world models over which predictions are made.

## Acknowledgement

For helpful conversation, we thank Bria Long, Brian Scholl, Felix Chang, Justin Halberda, Hongjing Lu, and the members of the Harvard Vision Sciences Laboratory. We also thank Anika Vaishampayan and Jeff Chang for their help in data collection. This work was supported by an NSF CAREER BCS-1942438.

## Author Contribution

Y.-C. Chen and T. Konkle designed the research and wrote the manuscript with input from A. Deza. Y.-C. Chen and A. Deza created the stimuli. Y.-C. Chen conducted the experiments and analyzed the data with input from T. Konkle.

## Open Practice Statement

The supplementary file available online with this paper contains the preregistration, materials, and raw data for each experiment, which can be found at https://osf.io/pqvsr/?view_only=c36c5cbad60f4228a37230590db4fde3.

## Appendix A: Texform Generation Procedure

### Image collection

Images of objects and animals were collected from various sources including stimuli from previous works (Long, Yu, & Konkle, 2018; Konkle & Caramazza, 2013; Konkle & Oliva, 2012) and Google images. We first removed the background of the images, and then cropped it to the square bounding box of the objects or animals. Only images with resolution higher than 512 px × 512 px after this step were included, and repetitions of objects from the same basic level category were replaced. This results in a total of 180 images, with 45 big objects, 45 small objects, 45 big animals, and 45 small animals.

### Normalization

All images were resized to 512 px × 512 px and converted to grayscale using the Rec. 601 Luma coding formula (red channel × 0.299 + green channel × 0.587 + blue channel × 0.114). They were then equalized across luminance and luminance histograms using the Spectrum, Histogram, and Intensity Normalization (SHINE) Toolbox (Willenbockel, Sadr, Fiset, Horne, Gosselin, & Tanaka, 2010). This step was done ignoring the background and without optimizing the structural similarity (SSIM) index (for details on the SSIM index, see Wang, Bovik, Sheikh, & Simoncelli, 2004). The images were then placed and centered on a gray (#828282) background of 640 px × 640 px.

### Texform Generation

The texform images were generated with an accelerated texform model (Deza, Chen, Long, & Konkle, 2019) modified from the method used in previous studies (e.g., Long, Yu, & Konkle, 2018): The preprocessed intact images were placed in a simulated visual periphery as an input image to a metamer model (Freeman & Simoncelli, 2011). Then, a metamer image was synthesized by iteratively coercing random noises to match the texture statistics of the input image for every overlapping simulated receptive field, in addition to roughly matching the structure given a low-pass residual of the input image. The procedure was run for 50 iterations using a variant of gradient descent, producing a final texform image, which is essentially a peripheral metamer of the intact image. The same intact image was passed into the model twice to generate a left and a right texform corresponding to the left and right visual periphery. This resulted in 180 left texforms, and 180 right texforms in total.

## Appendix B: Texform Images Recognizability Pretest

To assess the recognizability of the texform images, we used a similar free-guessing method used in a previous study (Long et al., 2018). Thirty-six subjects from Amazon Mechanical Turk (Mturk) participated in the recognition task. They viewed the texform images and guessed what they were. (For a discussion of this pool’s nature and reliability, see Crump, McDonnell, & Gureckis, 2013.) All subjects were in the U.S., had an MTurk task approval rate of at least 95%, and had previously completed at least 50 MTurk tasks.) Half of the subjects were shown the 180 left texforms, and the other half seen the 180 right texforms. The texforms were shown one by one and the subjects simply typed their guesses in a textbox without time constraint. The subjects were instructed to always give one answer only (e.g., avoid answers like “A or B”, “I don’t know”, or leaving the box blank).

Next, another 9 Mturk subjects (with the same Mturk qualifications) evaluated the guesses for the 360 texforms made from 180 original images. The original images were divided into 9 sets of 20 original images, and each subject viewed two sets of the original images (40 original images in total, where 10 from each of the 4 types: big objects, small objects, big animals, and small animals), with each set of the images viewed by two subjects. They viewed the images one by one along with 18 guesses for the corresponding texforms from subjects in the recognition task. The subjects in the evaluation task were never shown any texforms. For each subject, one set of the images were paired with the 18 guesses came from the 18 subjects who viewed the left texforms, and the other set were paired with the 18 guesses came from the 18 subjects who viewed the right texforms. Each set of the stimuli were evaluated by two subjects, with one subject evaluating the guesses for their left texforms, and the other subject evaluating the guess for their right texforms. The subjects were instructed to judge whether each guess could be “used to correctly describe” the original image. They were asked to consider guesses correct as long as the guesses were descriptions of something of similar real-world sizes and shapes from the correct answers. The guesses were spell checked and corrected using Microsoft Word before being shown to the subjects. We left the few guesses that were incorrectly and ambiguously spelled as is, since none of the likely interpretations of these guesses could influence the results. The count of guesses that were graded as correct yielded a recognizability score (ranging from 0 to 18) for each texform. We selected a final set of 40 pairs of intact and texform images (20 big items and 20 small items, half depicting animals and half depicting inanimate objects), where the items received a recognizability score of no more than 3 out of 18. The recognizability scores for the selected set can be found at https://osf.io/pqvsr/?view_only=c36c5cbad60f4228a37230590db4fde3.

## Appendix C: Individual Data Visualizations

**Figure A1.**
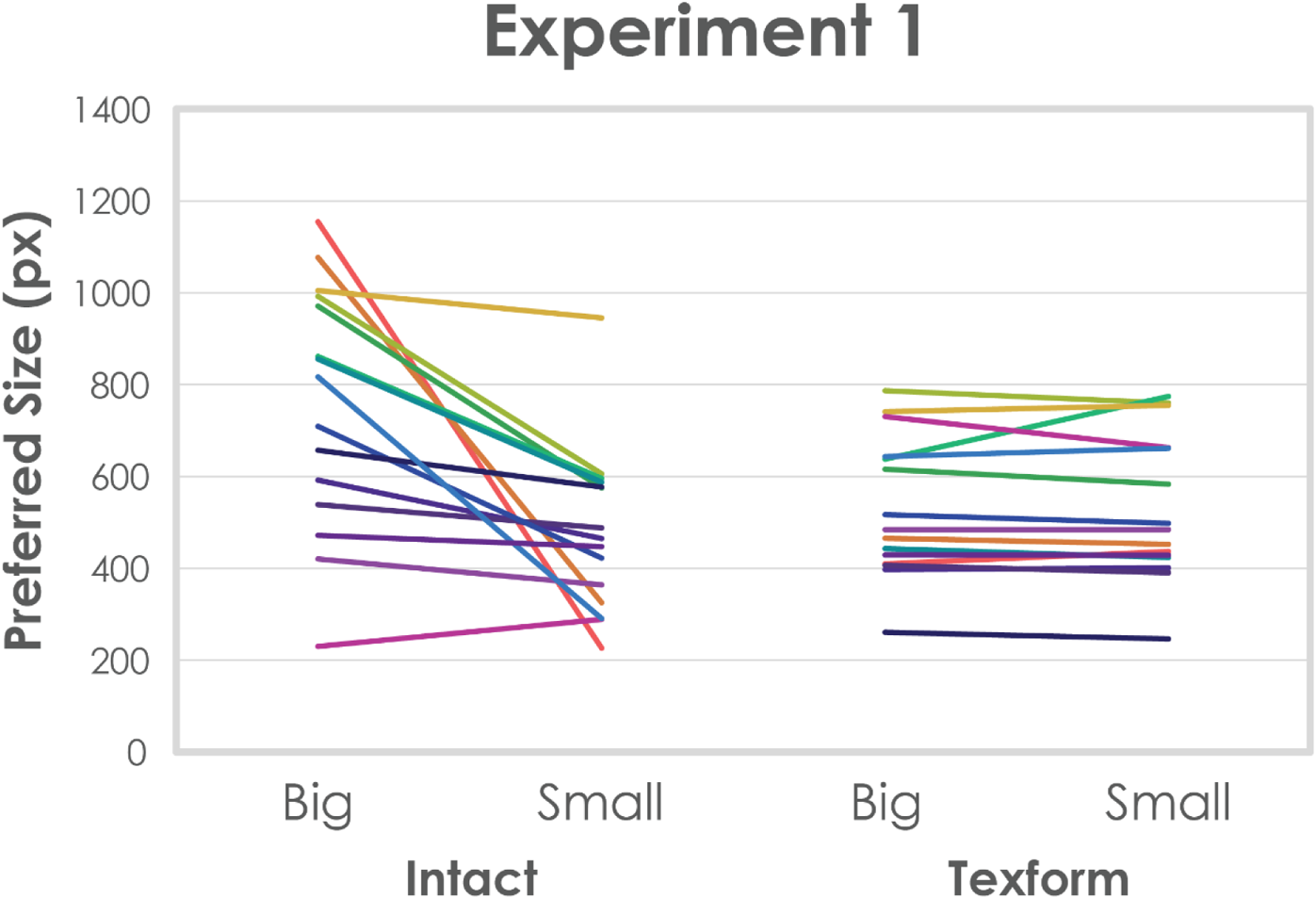
Each subject’s data from Experiment 1 are depicted with two lines of the same color (for intact and texform conditions respectively), with the selected size averaged over items plotted on the y-axis and the experimental conditions plotted on the x-axis.

**Figure A2.**
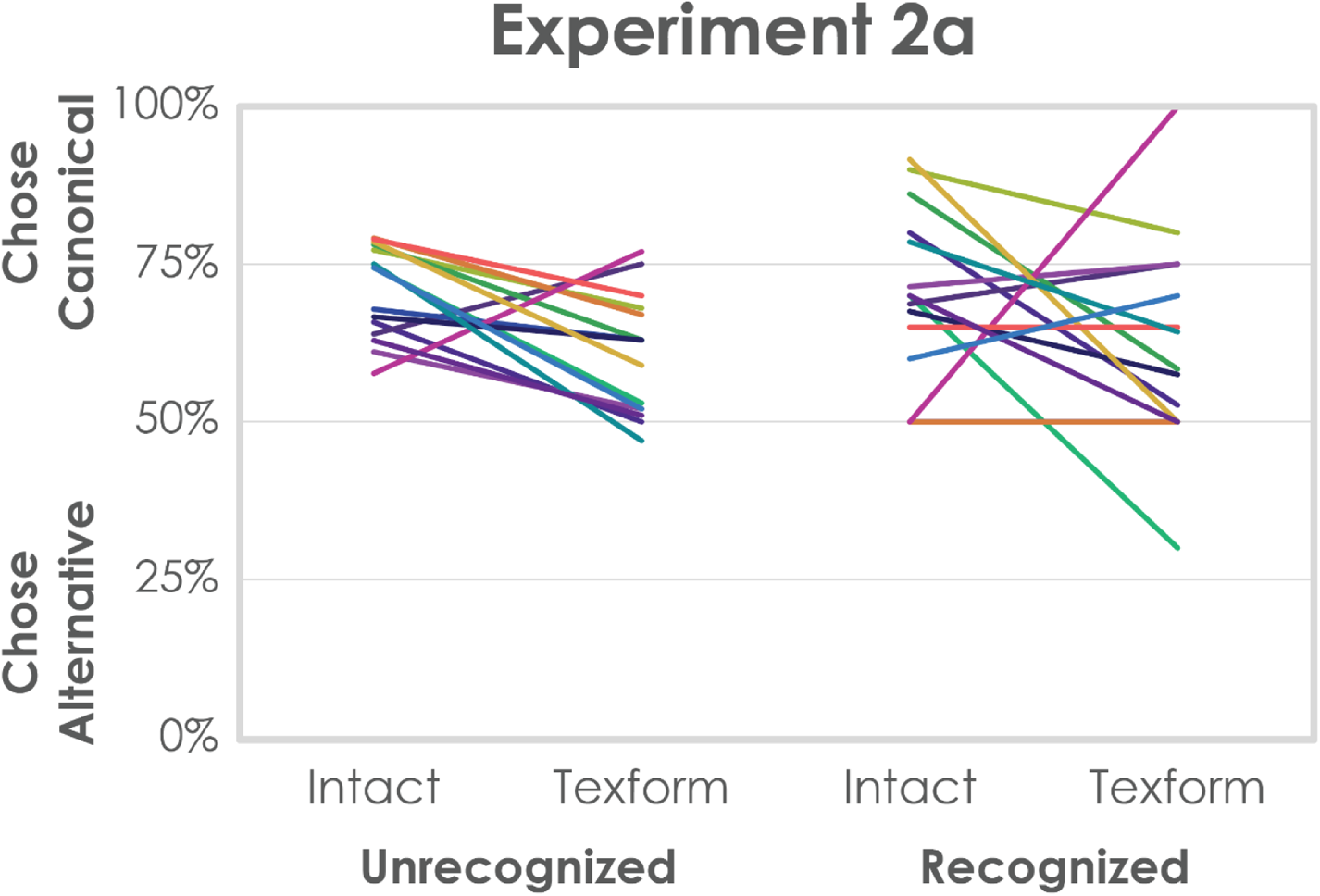
Each subject’s data from the choice task in Experiment 2a are depicted with two lines of the same color (for unrecognized and recognized items respectively), with percentage choosing canonical size plotted on the y-axis and the experimental conditions plotted on the x-axis.

**Figure A3.**
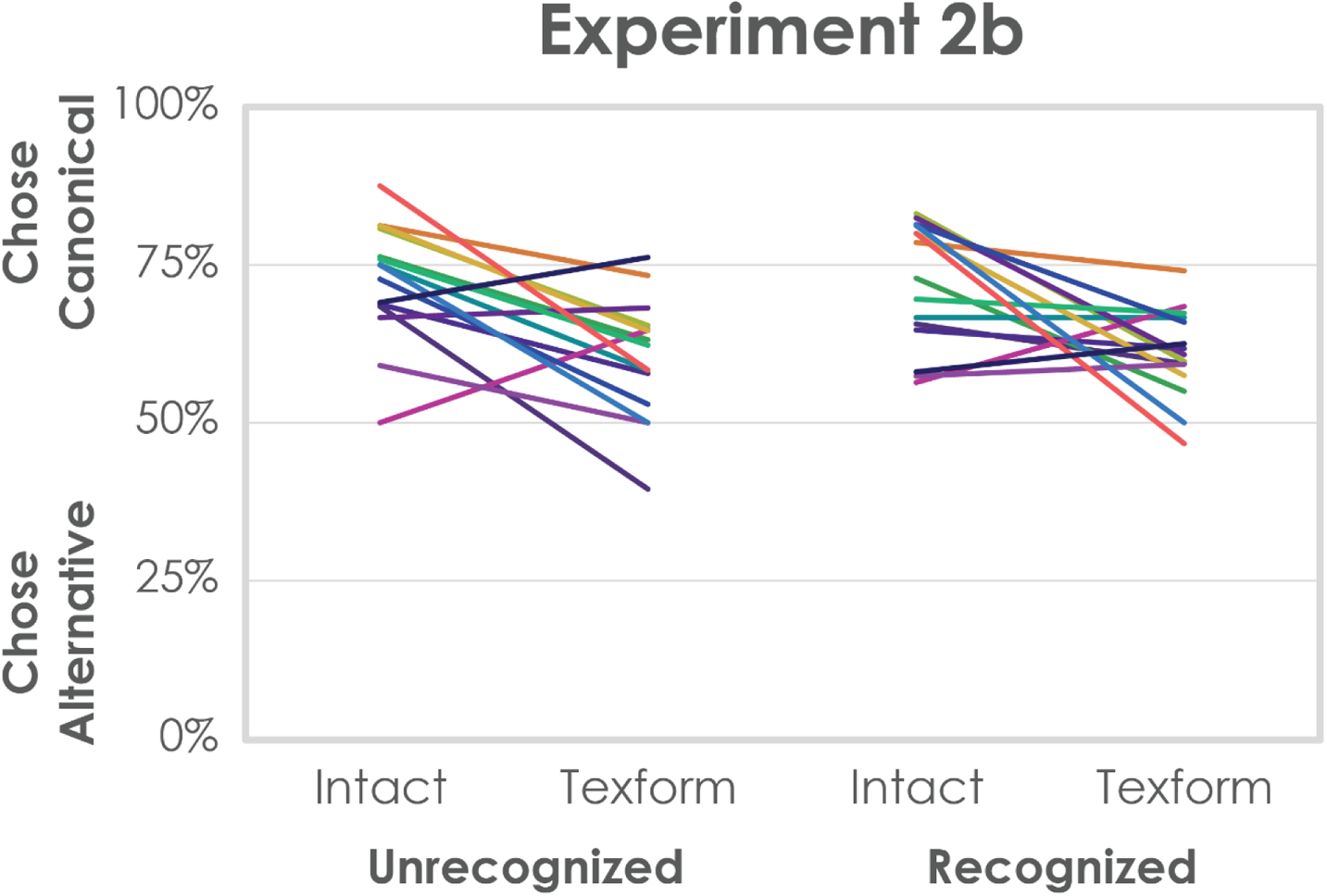
Each subject’s data from the choice task in Experiment 2b are depicted with two lines of the same color (for unrecognized and recognized items respectively), with percentage choosing canonical size plotted on the y-axis and the experimental conditions plotted on the x-axis.

**Figure A4.**
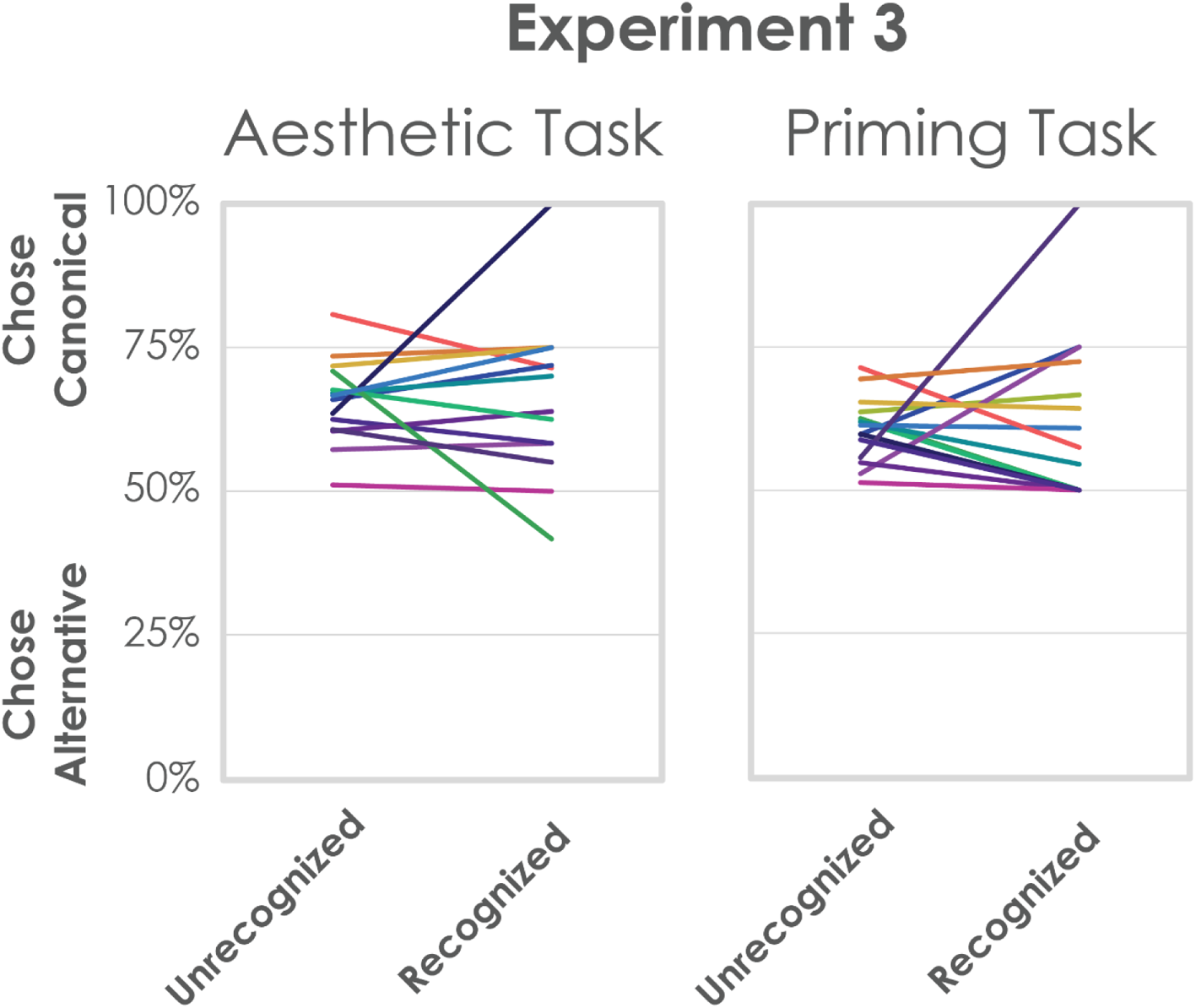
Each subject’s data from the choice task in Experiment 3 are depicted with a line, separately for aesthetic task and priming task conditions, with percentage choosing canonical size plotted on the y-axis and the recognition status plotted on the x-axis.

## Notes

1 For preregistration of Experiment 1, visit https://aspredicted.org/at3v7.pdf. The only deviation of experiment details from the preregistration is that the block order was not counter-balanced but alternated before subject exclusions.

2 For preregistration of Experiment 2a, visit https://aspredicted.org/pf87y.pdf. The only deviation of experiment details from the preregistration is that the block order was not counter-balanced but alternated before subject exclusions.

3 In their generation procedure the intact images were first scaled down to 180px × 180px and embedded in a 640px × 640px gray background. This image served as the input to generate the texform. After the synthesis, 192px × 192px area centered at where the input images were embedded was cropped and rescaled back to 440px × 440px. Finally, the four edges were gradually faded into the background color.

4 The preregistration specified 30 items as the criterion; however, this ended up being too strict and excluded most of the subjects. We thus decided on 12 items in Experiment 2a and replicated the results in Experiment 2b with this new criterion.

5 These analyses were done collapsing the animal/object factor as planned and preregistered. The results remained the same if we analyzed the animals and the objects separately (Experiment 2a intact block, animals: 70%, *SD*=10%, *t*(14)=7.47, *p*<.001, Cohen’s *d*=1.93, 14 out of 15 subjects, *p*=.001; objects: 72%, *SD*=8%, *t*(14)=10.28, *p*<.001, Cohen’s *d*=2.65, 15 out of 15 subjects, *p*<.001; texform block, animals: 60%, *SD*=8%, *t*(14)=4.80, *p*<.001, Cohen’s *d*=1.24, 12 out of 15 subjects, *p*=.035; objects: 61%, *SD*=13%, *t*(14)=3.52, *p*=.003, Cohen’s *d*=0.91, 12 out of 15 subjects, *p*=.035; Experiment 2b intact block, animals: 68%, *SD*=13%, *t*(14)=5.31, *p*<.001, Cohen’s *d*=1.37, 13 out of 15 subjects, *p*=.007; objects: 77%, *SD*=12%, *t*(14)=9.05, *p*<.001, Cohen’s *d*=2.34, 14 out of 15 subjects, *p*=.001; texform block, animals: 60%, *SD*=12%, *t*(14)=3.00, *p*=.009, Cohen’s *d*=0.78, 9 out of 15 subjects, *p*=.607; objects: 60%, *SD*=10%, *t*(14)=3.81, *p*=.002, Cohen’s *d*=0.98, 11 out of 15 subjects, *p*=.118).

6 The general aesthetic preference for curvatures in abstract shapes, objects, and interior designs have been well established in the studies cited here and beyond. However, it is important to distinguish the general preference from the potential role of curviness in the present context. Liking curvatures itself cannot explain the viewing size preferences, as we would only predict a general preference toward smaller objects that contain more curvy features regardless of viewing sizes.

7 This may initially appear to predict the opposite pattern in the viewing size preferences investigated here, but careful examinations in past studies have resolved this apparent conflict: The canonical viewing size of an object (on a screen) grow logarithmically with the real-world size of the object (Konkle & Oliva, 2011). At the same time, for an object with a fixed real-world size, the physical relationship between viewing size and viewing distance is linear. Thus, as real-world size of an object increases, the canonical viewing distance (varying with the canonical viewing size) increases as well (although by a smaller rate), predicting that we prefer to view smaller objects closer and bigger object farther.

## Notes

https://doi.org/10.17605/OSF.IO/PQVSR

### Competing Interest Statement

The authors have declared no competing interest.

### Summary of Updates

New experiment added.

